# Short-interval fires and vegetation change in southern California

**DOI:** 10.1101/2021.05.08.443193

**Authors:** Stephanie M. Lucero, Nathan C. Emery, Carla M. D’Antonio

## Abstract

**Questions:** In southern California, shortened fire return intervals may contribute to a decrease in native chaparral shrub presence and an increase in non-native annual grass presence. To test the hypothesis that short-fire return intervals promote a loss in shrub cover, we examined the contribution of single short-interval fires and abiotic conditions on the change of shrub cover within Ventura and Los Angeles counties. Through evaluating pre- and post-fire historical aerial images, we answered the following questions, 1) How has vegetation type cover changed after repeat fires? and 2) What landscape variables contribute the most to the observed change?

**Location:** Ventura County and Los Angeles County, California, USA.

**Methods:** We assessed the impact of a single short-interval fire by comparing vegetation recovery in adjacent once- and twice-burned fire burn polygons (long- and short-interval respectively). Pixel plots were examined within each polygon and vegetation cover was classified to vegetation type. We determined the best predictor of vegetation type cover with a linear mixed effects model comparison using Akaike Information Criterion.

**Results:** Pre-fire and post-fire community type cover was highly correlated. Burn interval was the best predictor of tree cover change (lower cover in twice-burned pixel plots). Aspect was the best predictor of sage scrub cover change (greater cover on north-facing aspects). Years since fire was the best predictor of chaparral cover change (positive correlation) and sage scrub cover change (negative correlation). Conversion of chaparral to sage scrub cover was more likely to occur than conversion of chaparral to annual grass cover.

**Conclusions:** Our study did not find extensive evidence of a decrease in chaparral shrub cover due to a single short-interval fire. Instead, post-fire cover was highly correlated with pre-fire cover. Chaparral recovery, however, was dynamic suggesting that stand recovery may be strongly influenced by local scale conditions and processes.

## Introduction

Fire-adapted ecosystems around the world are experiencing significant changes to the historical fire regime due to human influences (Piñol et al., 1998, Gillett et al., 2004, Syphard et al., 2007b, Nowacki and Abrams, 2008). The interval time between fire events (fire return interval), is one of the most direct way humans can alter a fire regime. Fire suppression has lengthened the interval time between fires in the northern Rockies (Barrett and Arno, 1982), in western Washington (Everett et al., 2000), and in the Sierra Nevada mountains (McKelvey et al., 1996) leading to with fire-adapted tree species being replaced by fire-sensitive, shade-tolerant species. A shorter fire interval can also put fire-adapted ecosystems at risk. In southern California, a shorter fire return interval is considered one of the leading drivers of native shrubs being replaced by non-native annual grasses (Keeley, 2000, Haidinger and Keeley, 1993, Zedler et al., 1983).

In southern California chaparral ecosystems, the historical fire return interval ranges from 30-90 years (Keeley et al., 2004, Van de Water and Safford, 2011) and in some locations may be as long as 150 years (Syphard et al., 2006). In this Mediterranean-type climate region, fires are typically crown fires (Hanes, 1971) occurring in the summer and fall (Beyers and Wakeman, 2000). Post fire, these stands generally return to pre-fire canopy cover within the first decade (Hope et al., 2007, Peterson and Stow, 2003) and to pre-fire stand structure within the second decade following fire (Hanes, 1971, Schlesinger and Gill, 1978).

In contrast, the current fire return interval can be much shorter due to an expansion of the wildland-urban-interface (Syphard et al., 2007a) and increased year-round anthropogenic ignitions caused by increasing human populations (Keeley and Fotheringham, 2001). Climate change is also expected to shorten mean fire return intervals as temperatures in southern California become warmer and precipitation more variable, thus increasing the risk of ignition (Krawchuk and Moritz, 2012, Polade et al., 2017). Furthermore, with the introduction of non-native annual grasses, chaparral communities could be driven toward a new successional trajectory leading to the extirpation of native shrub species and the expansion of non-native annual grasses (Brooks et al., 2004, Keeley and Brennan, 2012, D’Antonio and Vitousek, 1992).

A shortened fire interval has been recognized as one of the main causes of chaparral conversion to non-native annual grasses (Keeley, 2000, Syphard et al., 2019b). This is because many chaparral species require five-to-ten years to reach reproductive maturity (Zammit and Zedler, 1993) and at least 15 years to establish a robust seedbank (Syphard et al., 2018b, Park et al., 2018, Keeley, 2000). This is especially true of obligate seeding species, which are not fire resistant and germinate from seed following fire (Keeley, 1991). Extirpation of obligate seeding species and an increase in non-native species have been observed in the field when a second fire occurred in short succession (Zedler et al., 1983, Haidinger and Keeley, 1993, Keeley and Brennan, 2012, Jacobsen et al., 2004).

A single short-interval fire, however, does not always lead to chaparral conversion. Previous work with remotely sensed data suggest that water availability (Park et al., 2018, Syphard et al., 2019a), elevation (Meng et al., 2014), and mean annual temperature (Storey et al., 2021) explain more variation among chaparral stand recovery (although also see Syphard et al. (2019b)). These studies utilized imagery with 30-m spatial resolution and/or datasets with a 30-m scale.

There exists a research gap between in-person observations following a single fire event and landscape-scale observations across multiple fire events. The goal of this study was to address this limitation by using high resolution (1-meter) historical aerial imagery to observe detailed vegetation regrowth across the landscape and across multiple historical- and short-interval fire events.

In this study, we measured the difference in vegetation type cover following once- or twice-burned fire polygons (long- and short-interval fires, respectively) using historical aerial photographs ranging from 1956 and 2003 across two counties where, if chaparral conversion were occurring, we should be able to detect chaparral loss and decline. Through evaluation of pre- and post-fire images, we asked 1) How has vegetation type cover (i.e., chaparral, sage scrub, grass, tree) changed after repeat fires? and 2) What landscape variables contribute to the observed changes in vegetation cover?

## Methods

### Mapping the occurrence of short-interval fires

Fire history data (1879-2009) were acquired from the Fire and Resources Assessment Program (FRAP) database (CALFIRE, www.fire.ca.gov), reporting fires ≥ 4 hectares. The FRAP database was used because we required spatially explicit polygons for our analysis (although see Syphard and Keeley (2016) for the limitations of using the FRAP database). The shapefile was clipped in ArcMap (ArcGIS 10.1) to select for wildfires that fell within the study area of Ventura and Los Angeles counties and then processed to create an Interval Wildfire Occurrence map (Figure 1), each polygon having its own unique wildfire history based on original wildfire perimeters.

**Figure 1.**
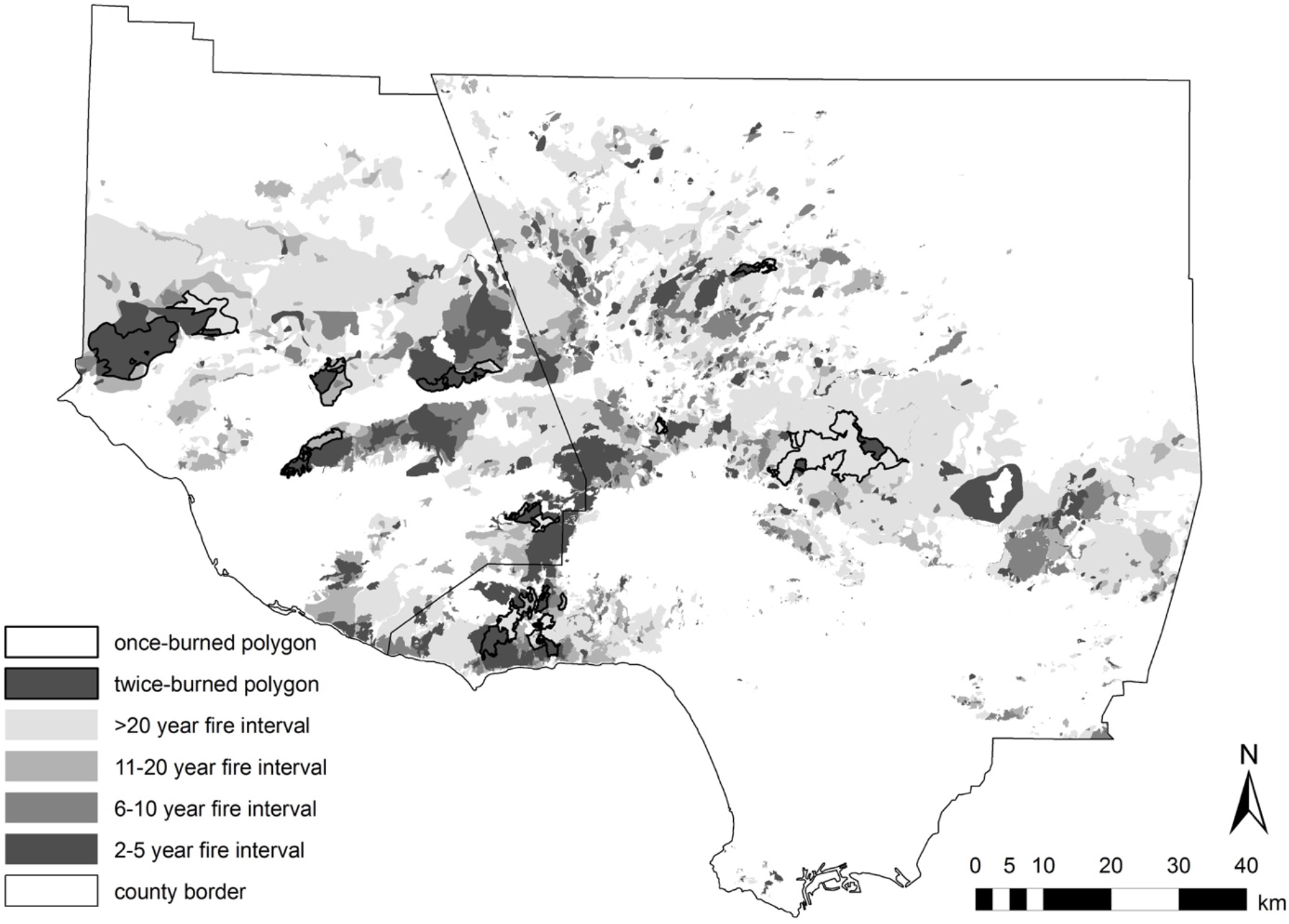
Minimum fire intervals as reported by CalFire (frap.fire.ca.gov) for Ventura and Los Angeles counties from 1878 to 2009. Twelve sites where one polygon burned twice within five years (twice-burned, black) and an adjacent polygon burned once within the same five-year period (once-burned, white).

The attribute table of the Interval Wildfire Occurrence map was exported to MS Excel (Microsoft 2011) and new metrics such as “Number of Fires” and “Minimum Fire Interval” were calculated (Appendix 1). Wildfire perimeters were filtered to eliminate single wildfires that were reported by multiple agencies, for example if a polygon had multiple “Fire Alarm Dates” in the same year (Jacobsen et al., 2004). The modified data table was finally joined back to the merged shapefile in ArcMap.

### Selection of twelve paired sites in Ventura and Los Angeles Counties

Ventura and Los Angeles Counties are ideal for determining the effects of a short-interval wildfire because the region is highly vulnerable to fire during the dry months (typically July-October) when Santa Ana wind conditions promote fast spreading wildfires (Hughes and Hall, 2010). In addition, the number of short-interval wildfires is predicted to increase as the population of southern California continues to grow (Keeley and Fotheringham, 2001, Myers and Pitkin, 2013).

Study sites were selected from the Interval Wildfire Occurrence map (Figure 1). We defined a “short-interval fire” as a fire return interval of five or fewer years. To quantify the effects of a single short-interval fire on vegetation, we selected polygons that experienced two wildfires within a five-year period that had an adjacent polygon that experienced only one wildfire within the same five-year period. All polygons had to be at least ≥0.5 km^2^ to allow for sufficient subsampling and analysis. Polygons that experienced a short-interval fire were considered “twice-burned” polygons (e.g., burned in 1962 and 1967) and adjacent polygons that only experienced the latter fire (e.g., 1967) were considered “once-burned” polygons. Selecting paired polygons that burned in the second fire, allowed for the polygons to experience the same number of post-fire recovery years. Long-interval polygons had, on average, 26.6 ± 5.2 (standard error mean) years of regrowth since the last wildfire and two of the twelve polygons (Site 3 and Site 9) had no prior record of wildfire since the early 1900s. For two sites (Site 4 and Site 6), long-interval polygons burned in the first wildfire year (instead of the second wildfire) and were analyzed in images ≥19 years post-fire. This exception was allowed assuming any difference in vegetation cover between long-interval and short-interval polygons would be negligible after ≥19 years of regrowth (Zammit and Zedler, 1993).

In addition to burn history, sites were selected to represent vegetation variation within the two counties and along a moisture gradient determined by their distance from the coast. Sites closer to the coast are generally more mesic and sites farther inland are generally more arid (Franklin, 1998). Water availability influences chaparral and sage scrub community composition and extent (Mooney and Parsons, 1973, Poole and Miller, 1981). Sites were selected to represent the vegetation communities within Los Padres National Forest in Ventura County and within the Angeles National Forest and the Santa Monica Mountain National Recreation Area in Los Angeles County.

### Selecting aerial photographs

Historical aerial photographs (HAPs) were acquired from the Map and Imagery Laboratory (MIL) at the University of California, Santa Barbara in 2011 to 2014 (www.library.ucsb.edu/mil). HAPs were chosen between 1952 and 2009 for corresponding wildfires spanning 1956 to 2003. Pre-fire HAPs were selected as close to before the first wildfire as possible to record initial vegetation cover and post-fire HAPs were selected ≥6 years following the second wildfire to capture maximal vegetation cover without encountering a third wildfire (Appendix 1). Vegetation communities were assumed to return to pre-fire canopy cover within six years following wildfire (Muller et al., 1968, Schlesinger and Gill, 1978). Seasonality of images was not controlled for under the assumption that mature communities appear distinguishable year-round (i.e., chaparral appears darker with a closed canopy year-round; grass appears lighter and has no visible canopy structure year-round). Final HAP selection was based on photograph availability and adherence to fire criteria.

### Georectifying aerial photographs

To compare pre- and post-fire vegetation cover on a pixel-by-pixel basis, all HAPs were georectified to the same base image. Grayscale, 2009, one-meter spatial resolution, digital orthophoto quarter quads (DOQQ) of Ventura or Los Angeles County, collected by the United States Geological Survey, were used as the base image. Temporally stable objects such as large shrubs or trees, rock outcrops, and crests and troughs of the mountainous landscape were used as registration points (RPs). Dirt roads and permanent structures were also used, although these more permanent features were rare in the HAPs due to the remoteness of the polygons. Because the terrain of the HAPs was mountainous and highly variable, RPs were placed at a high density to increase warping accuracy. Each HAP was then warped using triangulation and pixels were resampled to the nearest neighbor, creating a georectified HAP with one-meter spatial resolution.

Georectified HAPs (gHAPs) were then mosaicked to minimize edge distortion and increase spatial accuracy for vegetation analysis. Mosaicked gHAPs covered the entire long-interval and short-interval polygon of a site under pre-fire and post-fire conditions. Only two sites (Sites 2 and 6) were not georectified across their entire long-interval polygon due to a lack of available HAPs and/or their extensive size and instead an equivalent or greater area to the short-interval polygon was georectified.

Mosaicked gHAPs were validated for their spatial accuracy by identifying 40-100 RPs corresponding to the 2009 DOQQ base map. Validation RPs had a final root mean square error of ten pixels (i.e., ten meters) or less.

### Plot selection on north and south aspects within long- and short-interval polygons

Aspect is a large influencer of vegetation type cover (Hanes, 1971) with south facing aspects receiving more solar radiation than north facing aspects. Random points were generated within each site’s pre-fire gHAP and approximately eight points in the long-interval polygon and eight points in the short-interval polygon were selected for vegetation analysis. At each point, a 50 × 50 pixel plot (50 × 50 m) was established. Pixel plots (hereafter “plots”) were distributed between north and south facing aspects (north: 0.0° to 67.5° or 292.5° to 360°; south: 112.5° to 247.5°) to account for differences in solar irradiance (northern aspects receive less solar irradiance than southern aspects). Plots were shifted if needed to ensure they did not overlap a mountain ridge or valley and to fit entirely on one aspect. Aspect was verified with 30-meter USGS Digital Elevation Model (DEM) data and/or visually with Google Earth (Google Earth Pro 7.3.0.3832). All pre-fire plots were replicated in the post-fire mosaicked gHAP to capture vegetation regrowth at the same location.

In total, 198 plots were analyzed with 99 plots in long-interval polygons and 99 plots in short-interval polygons. One hundred and four plots had northern aspects and 94 plots had southern aspects. Plots were considered independent after including site as a covariate in plot level analysis and found no significant influence on plot level results.

Prescribed burns included in the FRAP database were reviewed and only two of the 198 plots overlapped with a prescribed burn. These two plots were removed from the analysis.

### Quantifying vegetation type cover within plots

To quantify vegetation type cover at each plot, the “dot grid” method was used (Floyd and Anderson, 1982, Dublin, 1991). A ten-by-ten grid (100 points) was overlaid on each plot with a spacing of five pixels (five meters) between each point. Vegetation cover was classified to vegetation type: chaparral, grass, sage scrub or tree. All grass cover was assumed to be non-native dominated based on the 1930’s Wieslander Maps and the 2001 USDA California Vegetation map. For classification consistency, all sites were examined twice to account for initial training and improvement in classification over time.

To improve classification accuracy, solar zenith was considered to account for shadows and Google Earth was referenced for vegetation type cover and seasonal changes (available years: 1990-2015). The authors traveled to six of the twelve sites to confirm that site cover approximated vegetation type cover observed in the HAPs (e.g., a matrix of chaparral, sage scrub, and grass in the HAPs were a matrix of vegetation in the field). However, verification of the HAPs was infeasible due to many of the images being decades old.

Percent cover (%) was quantified by tallying the number of points classified within each vegetation type (100 points = 100% cover). Pre-fire vegetation cover was subtracted from post-fire vegetation cover to quantify the amount of vegetation type change within each plot.

### Datasets for abiotic variables

Aspect was calculated from USGS digital elevation models (DEMs) with a 30 × 30 meter horizontal resolution and a one-meter vertical resolution. Distance from the coast was calculated in ArcMap by determining the centroid point of each polygon and measuring the shortest direct distance to the coastline (Appendix 1).

Moisture availability following fire, influences seedling survival and thus eventual vegetation type cover (Pratt et al., 2014, Venturas et al., 2016). It was calculated from PRISM (http://www.prism.oregonstate.edu/explorer/) averaging the annual precipitation during the first five years of regrowth following the latter wildfire.

### Statistical analysis

Statistical analysis was conducted using RStudio (RStudio, Inc. version 0.98.1103) and R version 3.3.2 (http://www.rstudio.com/) and were either run at the plot level (e.g., 99 once-burned and 99 twice-burned plots) or at the polygon level (e.g., 12 once-burned polygons and 12 twice-burned polygons). Polygon values were calculated as the mean of their plot values.

As post-fire vegetation type cover at the plot level was highly correlated with pre-fire vegetation type cover (Figure 2), the residuals of a linear regression for pre- and post-fire vegetation types were derived to account for pre-existing plant communities. A linear mixed-effects model of plot data (N = 198) was run for each vegetation type with site as a random effect to reduce the effects of spatial autocorrelation. The models to predict the residuals were: burn interval, aspect, years after fire (when post-fire HAPs were taken), distance to coast, and the five-year average rainfall post-fire. In total, five models were run for each vegetation type. The Akaike Information Criterion (AIC) values of each model were compared within a vegetation type. The model(s) with the lowest AIC value were further investigated for each vegetation type. If the best model predictor was categorical, then an ANOVA was performed to determine statistical significance and if the model predictor was continuous, a linear regression was performed to determine significance.

**Figure 2.**
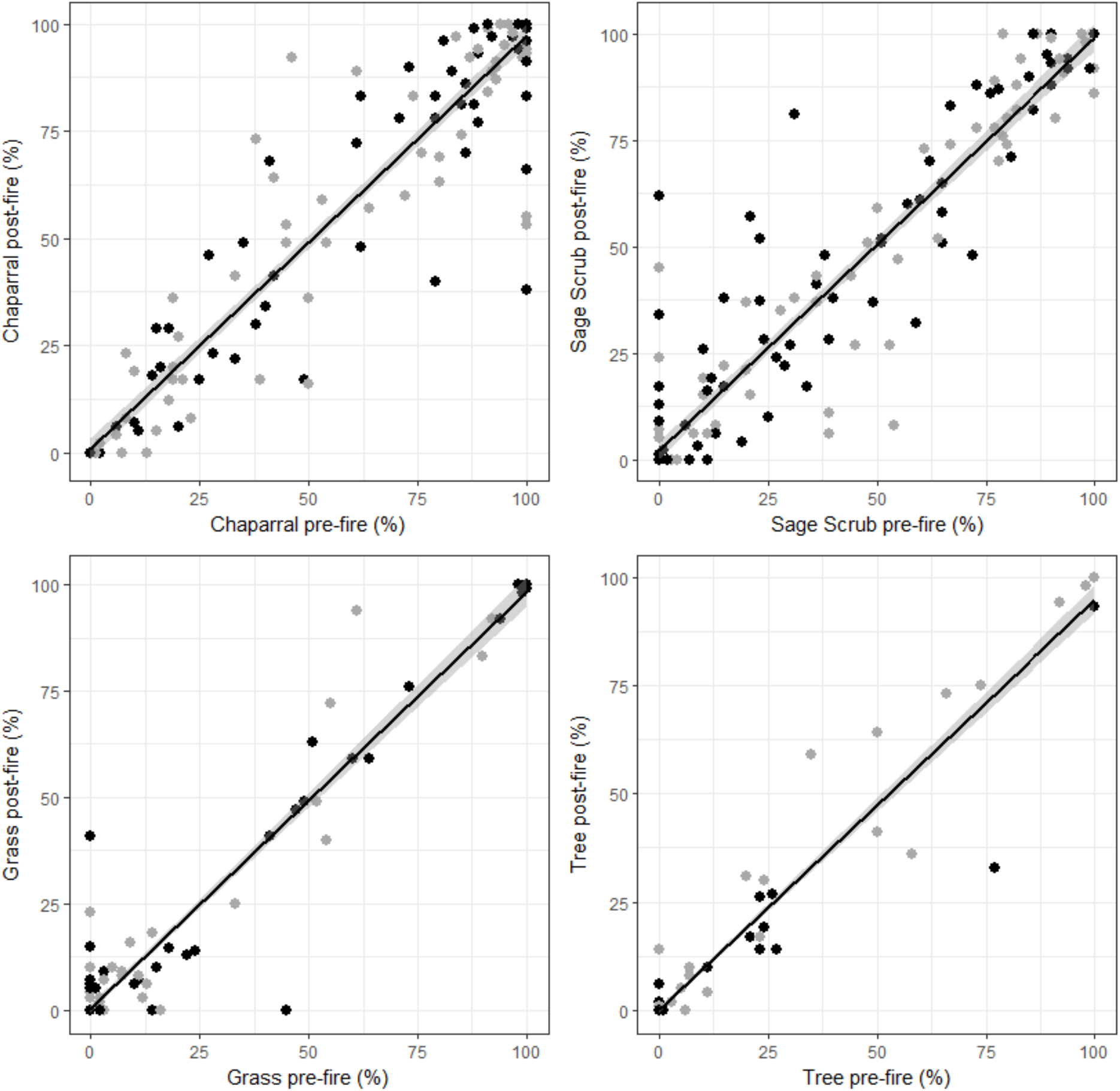
Linear regressions with confidence intervals of pre-fire and post-fire percent cover for all four vegetation type covers. Each point represents a once-burned (gray) or twice-burned (black) plot.

## Results

### Trends in vegetation type cover comparing pre-fire and post-fire conditions

For each vegetation cover type, post-fire conditions were highly correlated with pre-fire conditions (N = 198) (Figure 2). Linear regressions were significant for chaparral percent cover (*P* < 0.001, r^2^ = 0.93), sage scrub percent cover (*P* < 0.001, r^2^ = 0.91), grass percent cover (*P* < 0.001, r^2^ = 0.93), and tree percent cover (*P* < 0.001, r^2^ = 0.94).

### Linear mixed-effects model

When analyzing chaparral vegetation type residuals, the models with the lowest AIC values included aspect and years since fire (Table 1). An ANOVA of aspect yielded a trend towards a difference between north and south facing aspects (*P* = 0.083, F = 3.05) with south-facing aspects showing greater decreases in cover than north-facing aspects (Figure 3, top right). Change in chaparral cover was positively correlated with longer recovery time post-fire (Figure 3, top left, *P* < 0.001, t = 3.507). For sage scrub residuals, the models with the lowest AIC were also aspect and years after fire. Plots on north-facing aspects had greater positive change in sage scrub percent cover than south-facing slopes (Figure 3, middle right, *P* = 0.034, F = 4.567). Change in sage scrub cover was negatively correlated with years after fire (Figure 3, middle left, *P* = 0.005, t = −2.823). The lowest AIC value for grass residuals was years after fire, however the difference in AIC value among models was very slight. A linear regression of the change in percent grass and years after fire was not significant (Figure 3, bottom left, *P* = 0.185, t = −1.332). For tree residuals, the best model was burn interval. Once-burned plots had more negative change in tree cover than short-interval plots (Figure 4, ANOVA, *P* = 0.030, F = 4.769). This pattern was still significant when the outlier site (Site 11, Plot 7: −39.97) was removed (ANOVA, *P* = 0.048, F = 3.97).

**Table 1.** Linear mixed-effects model results for all vegetation type covers at the plot level (N = 198). Each model had site as a random effect and was analyzed using the residuals of pre/post-fire vegetation percent cover.

| Vegetation Class | Model AIC |  |  |  |  |
| --- | --- | --- | --- | --- | --- |
|  | Burn Interval | Aspect | Years After Fire | Distance to Coast | Five-year Average Rainfall |
| chaparral | 1519.907 | 1515.694* | 1514.127* | 1519.806 | 1519.095 |
| sage scrub | 1525.744 | 1522.241* | 1522.682* | 1527.828 | 1527.542 |
| grass | 1296.388 | 1296.534 | 1295.17* | 1296.466 | 1296.417 |
| tree | 1160.116* | 1164.776 | 1164.47 | 1164.837 | 1164.873 |

\* indicates best model(s)

**Figure 3.**
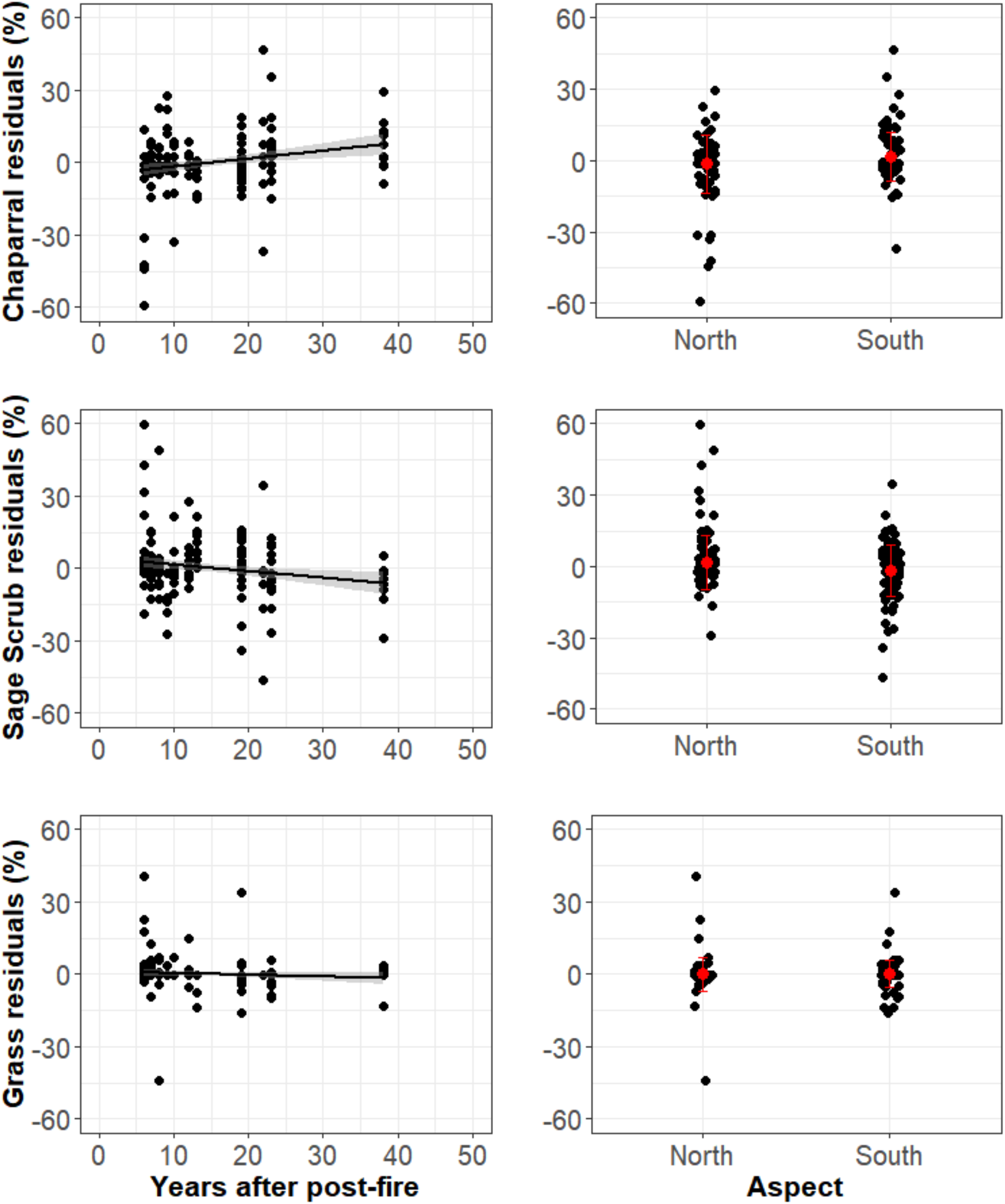
Linear regressions with confidence intervals of time since last fire with residuals (left side) for chaparral (*P* < 0.001***, r^2^ = 0.05), sage shrub (*P* = 0.005**, r^2^ = 0.03), and grass (*P* = 0.16, r^2^ = 0.005) cover. Distribution of the residuals (right side) of chaparral (ANOVA, *P* = 0.114), sage scrub (ANOVA, *P* = 0.046*), and grass (ANOVA, *P* = 0.886) cover by aspect.

**Figure 4.**
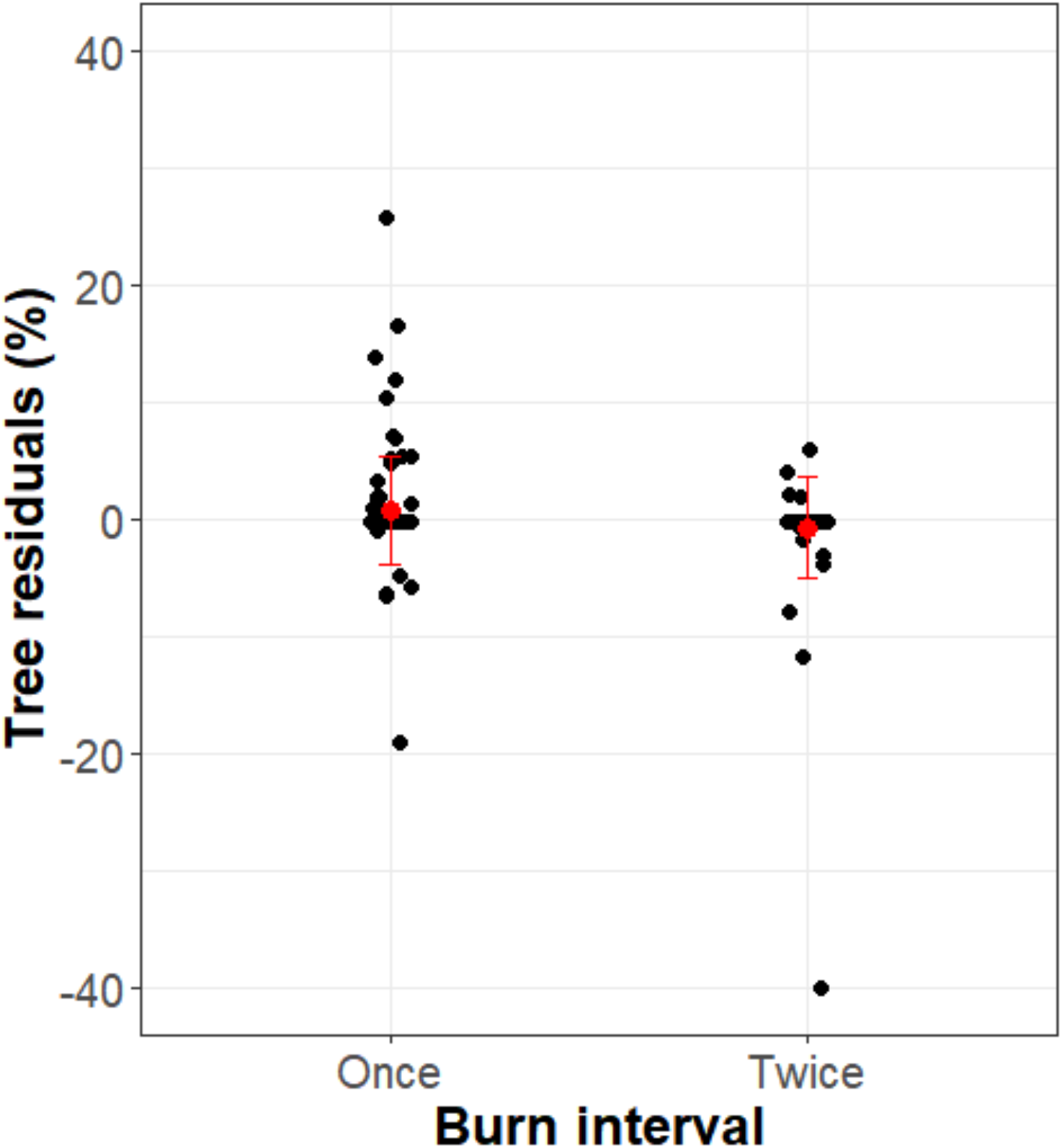
Distribution of residuals for tree cover (ANOVA, *P* = 0.03*) by burn interval. When the outlier of tree percent lost in the twice-burned burn interval is removed, the relationship is still significant (ANOVA, *P* value = 0.048*).

### Transition matrix

The majority of pixels showed no change in cover (Figure 5). The majority pixels located in once-burned plots returned to their pre-fire cover (chaparral: 86.3%; sage scrub: 80.4%; grass: 76.3%; tree: 74.0%). Pixels in twice-burned plots also often returned to their pre-fire cover (chaparral: 87.1%; sage scrub: 78.4%; grass: 70.0%; tree: 65.7%).

**Figure 5.**
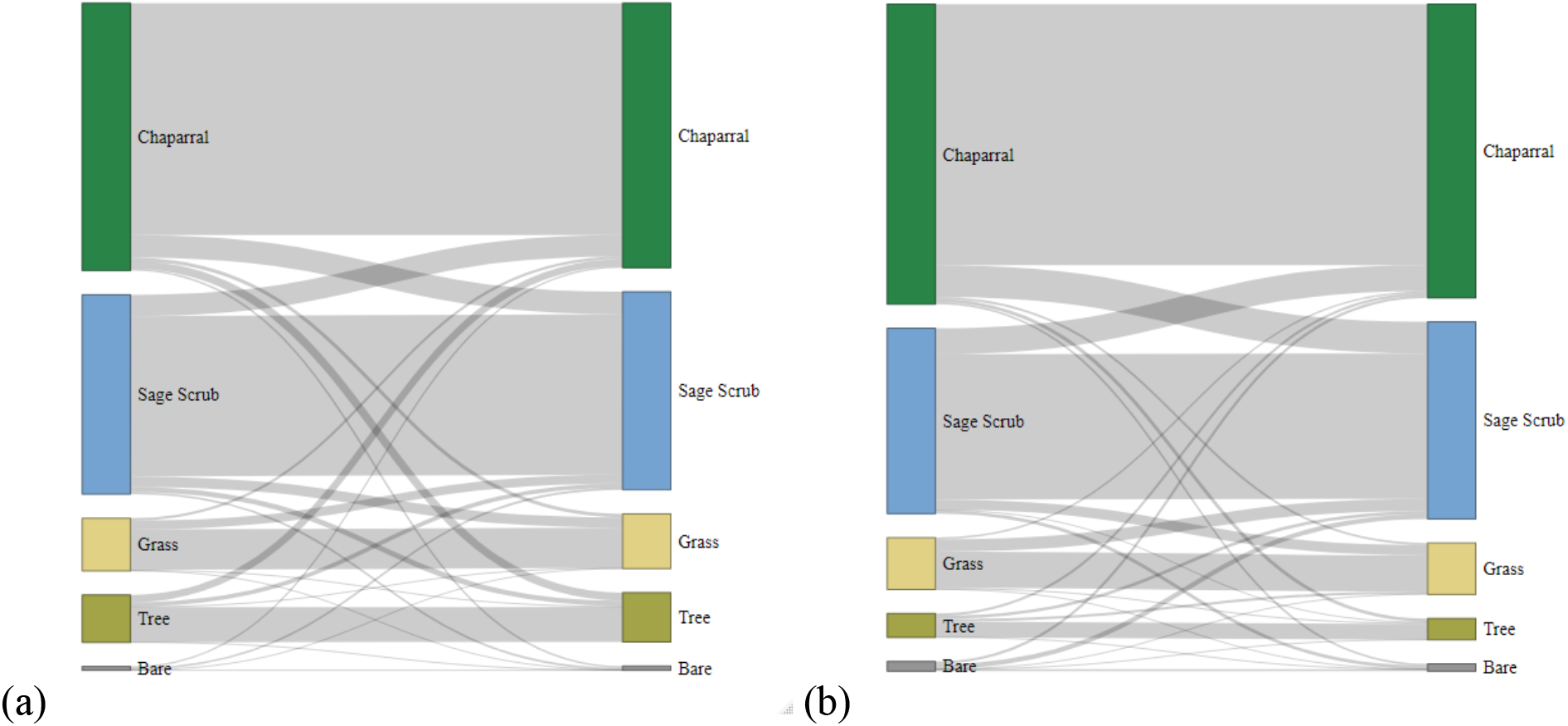
Transitions for all pixels showing pre-fire (left axis) and pos-tfire (right axis) cover within (a) once-burned plots and (b) twice-burned plots. Gray bands depict the proportion of pixels that remained or transitioned to another cover type.

The transition direction and the proportion of pixels that changed in cover were similar between once- and twice-burned locations. Chaparral pixels were more likely to convert to sage scrub than to grass (once-burned: 8.4% and 1.3% respectively; twice-burned: 10.5% and 0.7% respectively). Sage scrub pixels were more likely to convert to chaparral than to grass (once burned: 10.8% and 5.2% respectively; twice-burned: 14.0% and 5.5% respectively). Grass pixels that converted were most likely to convert to sage scrub cover (once-burned: 17.0%; twice-burned: 23.8%).

## Discussion

The results of this study found that a single short-interval fire, as quantified in twice-burned plots, did not lead to significant vegetation type change of chaparral, sage scrub or grass cover at the landscape scale. A short-interval fire was only a significant predictor of reduced vegetation cover for plots dominated by trees (Table 1). Instead, overall vegetation recovery trends were driven by the pre-existing cover (i.e., post-fire cover was significantly correlated with pre-fire cover) (Figure 2).

There were, however, a large number of plots that either increased or decreased in cover about the mean. Suggesting there is greater heterogeneity at the individual plot level than there is at the landscape level. Ninety-eight out of 198 chaparral plots and 91 out of 198 sage scrub plots showed an increase or decrease in woody cover following fire. This heterogeneity in canopy regrowth is consistent with previous findings (Syphard et al., 2018a, Park et al., 2018, Storey et al., 2021).

While we set out to observe if there was evidence of chaparral conversion to grass following a single short-interval fire, we found there was greater conversion from chaparral to sage scrub (10.5%) than chaparral to grass (0.6%). In comparison, sage scrub was more likely to convert to grass (5.5%). These interactions may be influenced by these communities’ physical distribution. Sage scrub communities are typically found below chaparral at lower elevations, which tend to be hotter and drier, and adjacent to urban areas, which tend to be dominated by annual grasses. Similar results were found by Syphard et al. (2018a, 2018b, 2006) and Meng et al. (2014).

Aspect was a significant driver of sage scrub cover (Figure 3, middle right) and a strong driver of chaparral cover (Figure 3, top right) with cover increasing on north-facing aspects and decreasing on south-facing aspects. This result was unexpected as north-facing aspects tend to have more mesic conditions and receive less solar radiation, conditions where chaparral typically outcompete sage scrub (Syphard et al., 2006). Miller et al. (1983) showed similar dry down periods (0-2 weeks) between north- and south-facing aspects, indicating soil moisture on opposing aspects may not be as different as originally expected.

There was strong evidence that chaparral cover increased on north-facing aspects and decreased on south-facing aspects (Figure 3, top right), which is consistent with previous research (Hanes, 1971, Keeley and Keeley, 1981) and other studies that found metrics of soil aridity to be positively correlated with chaparral loss (Syphard et al., 2019a) and grass presence (Park et al., 2018).

Another explanation as to why no significant difference was detected between north- and south-facing aspects could be due to the expansion of *Malosma laurina*, a tenacious facultative resprouter, on south-facing aspects, which can outcompete other species when water resources are limited (Thomas and Davis, 1989). Because *M. laurina* can reach similar heights and canopy densities as mixed chaparral stands, locations dominated by *M. laurina* would have been classified as “chaparral” in the HAPs.

While we were readily able to differentiate between vegetation cover types (e.g., grass, sage scrub, chaparral, tree), we were not able to identify vegetation to species, which meant we were unable to detect if there was a change in community composition or a change in the presence of obligate seeding, obligate resprouting, or facultative resprouting shrub species. These reproductive strategies are predicted to respond differentially to increased fire frequency (Keeley, 1991, Syphard et al., 2006, Franklin et al., 2004) and distribution can vary by water availability (Mooney and Parsons, 1973, Poole and Miller, 1981, Franklin, 1998, Meentemeyer and Moody, 2002).

In addition, no significant difference in chaparral cover on north- or south-facing aspects could be due to the method of data collection. Since values from the dot-grid method spanned 0-100%, plots with close to 100% chaparral cover pre-fire had little room to increase in chaparral cover post-fire. Within plots that had 100% chaparral cover pre-fire, a decrease in cover or “no change” was the only possible outcome. Indeed, average pre-fire chaparral cover on north-facing (more mesic) aspects was 72.19 ± 3.78% compared to only 26.33 ± 3.49% on south-facing (more arid) aspects. North-facing aspects had more chaparral cover to lose at the outset.

Calculating the relative change in vegetation cover was explored, however it inflated non-biologically important results (e.g., 1 pixel post-fire / 2 pixels pre-fire = decrease of 50%) and hid larger community changes (e.g., 60 pixels post-fire / 100 pixels pre-fire = decrease of 40%). For this reason, all changes in cover were calculated as their absolute value (e.g., 1 pixel post-fire – 2 pixels pre-fire = decrease by 1).

Chaparral and sage scrub cover showed strong yet opposing trends in succession following fire. Chaparral cover increased significantly with additional years after fire whereas sage scrub cover strongly declined with additional years after fire (Figure 3, left). This supports previous studies that suggest sage scrub can be successional to chaparral given enough time without fire (McPherson and Muller, 1967, Gray, 1983). This transition in canopy cover is consistent with the assumption that chaparral stands require five to thirty years to recover after wildfire (Hanes, 1971, Hope et al., 2007, Schlesinger and Gill, 1978) and subshrubs, common in sage scrub communities, decline as chaparral shrubs mature around them (Syphard et al., 2006). Thus, it could be possible that with enough fire-free years, chaparral cover could expand via the establishment of chaparral species within sage scrub stands.

Number of years post-fire and vegetation succession may also explain some of the vegetation change detected in Syphard et al. (2018a). Some locations observed to convert from shrub to grass cover may have been dominated by or included herbaceous cover in the process of recovering following fire and may not represent the mature community. The Day fire (2006), Zaca fire (2007), and La Brea fire (2009) occurred within seven years prior to the 2013 Landfire vegetation map, so it is within the window of expected recovery that some of these locations were still recovering and may eventually return to shrub cover absent of short-interval fires.

We hypothesized sites closer to the coast would be more mesic and sites farther from the coast would be more arid, leading to less vegetation conversion near the coast and more conversion inland. We did not, however, find distance to the coast to be a significant factor. This could be due to vegetation trends being more strongly correlated with moisture gradients driven by elevation than by distance from the coast. Indeed, Meng et al. (2014) and Syphard et al. (2019b) found elevation to be a strong driver of vegetation change with more chaparral conversion occurring at lower elevations. Analysis by Storey et al. (2019) included a more extensive range of chaparral stands in southern California and found vegetation change was greatest in the eastern portion of their study region and at higher elevations and was a result of drought and fire interactions.

The five-year average rainfall following the latter fire was also found to not be a significant driver of vegetation type change, which is in line with other studies (Storey et al., 2020, Meng et al., 2014), although see Storey et al. (2021). Storey et al. (2020) found climactic water deficit to be a stronger predictor of chaparral recovery compared to total precipitation and water conditions preceding fire were generally more predictive of recovery than conditions following fire. Finer temporal scale patterns of rainfall (e.g., light rainfall over a month vs. a one-day downpour, early winter vs. late winter) will also likely be important to consider in the seasons leading up to fire and the first year following fire. This will especially be true of locations that are already water limited (e.g., < 500 mm y^−1^).

While almost all polygons included in this study experienced additional fires before and after the years of analysis, multiple short-interval fires were not included in this analysis nor was the entire fire history (1878-2009) of a location. Storey et al. (2021) included additional fire history by comparing locations in southern California that burned once, twice, or three times within 25 years. They found no difference in shrub recovery between stands that burned once or twice but did find a decrease in shrub recovery when locations burned three times within 25 years. Investigating sites that experienced multiple short-interval fires will continue to be highly valuable as will investigating the impact of variability in years between fires (Zedler, 1995) on vegetation change.

Another factor that might have contributed to our results is the time frame of analysis (i.e., 1956-2003). Resilient shrub species may have already been selected for in locations that experienced multiple fires. Thus, any vegetation change that would have occurred due to a short-interval fire may have already occurred, prior to when imagery was first available.

While this study did not find a consistent increase in grass cover following a single short-interval fire, non-native annual grasses still pose a risk to native plant communities. When non-native annual grasses senesce they create highly ignitable, continuous fuel across the landscape that can carry fire into chaparral stands that otherwise would not have ignited (Keeley, 2000). An expanding wildland-urban-interface (Syphard et al., 2007b) and climate change (Krawchuk and Moritz, 2012) are also expected to create additional risks for southern California’s wildlands.

## Conclusion

The aim of this study was to identify how vegetation type cover changed in response to a single short-interval fire (defined as two fires within five years) and how environmental variables contributed to vegetation type change across the landscape. We did not find extensive evidence that short-interval fires promoted a landscape-scale loss of shrublands. Instead, we found chaparral cover only slightly declined in twice-burned plots. Tree cover was the only vegetation type to significantly decline following a single-short interval fire. Our data are consistent with the hypothesis that chaparral conversion to other vegetation types occurs slowly and is likely influenced by its abiotic environment. Aspect was the best predictor of differences in sage scrub cover while the number of years post-fire was the best predictor of chaparral and sage scrub stand regrowth. Post-fire and pre-fire cover, overall, were highly correlated for all vegetation cover types, although variation at the pixel plot level was apparent, suggesting that vegetation change occurring at the local scale may be influenced primarily by local scale processes. Therefore, caution is advised when managing fire-prone vegetation types as these landscapes can be highly heterogeneous and outcomes at one location may not reflect landscape level patterns.

## Acknowledgements

We are grateful to E. Burley, S. Suresh, C. Allred, T. Madden, and M. Plummer for their assistance processing aerial images; to S. Peterson for guidance in ENVI and ArcGIS; and to P. Dennison, M. Moritz, and R. Oono for their advice, guidance, and feedback on earlier versions this manuscript.

